# Insulin Receptor Deletion in S100a4-lineage cells accelerates age-related bone loss

**DOI:** 10.1101/445155

**Authors:** Valentina Studentsova, Emma Knapp, Alayna E. Loiselle

## Abstract

Type I and Type II Diabetes dramatically impair skeletal health. Altered Insulin Receptor (IR) signaling is a common feature of both diseases, and insulin has potent bone anabolic functions. Several previous studies have demonstrated that loss of IR in bone cells results in disrupted bone homeostasis during early post-natal growth. Here we have deleted IR in S100a4-lineage cells (IRcKO^S100a4^) and assessed the effects on bone homeostasis at both young (15 weeks) and older adult (48 weeks) mice. S100a4-cre has previously been shown to target the perichondrium during bone development, and here we show that S100a4 is expressed by adult trabecular and cortical bone cells, and that S100a4-Cre effectively targets adult bone, resulting in efficient deletion of IR. Deletion of IR in S100a4-lineage cells does effect initial bone acquisition or homeostasis with no changes in cortical, trabecular or mechanical properties at 15-weeks of age, relative to wild type (WT) littermates. However, by 48-weeks of age, IRcKO^S100a4^ mice display substantial declines in trabecular bone volume, bone volume fraction and torsional rigidity, relative to age-matched WT controls. This work establishes the utility of using S100a4-cre to target bone and demonstrates that IR in S100a4-lineage cells is required for maintenance of bone homeostasis in adult mice.

## Introduction

Both Type I and Type II Diabetes dramatically impair skeletal health. Type I Diabetes both accelerates the onset and increases the risk of osteopenia and osteoporosis [1-3], while Type II Diabetes can actually increase bone mineral density [4, 5]. However, in both cases, bone homeostasis is disrupted, including decreased bone quality (reviewed in [6]), resulting in a substantial increase in fracture risk [7, 8]. Moreover, in both clinical and pre-clinical models impaired healing following bone fracture is observed in Type I and Type II Diabetics [9-12]. Despite their different pathophysiologies, disruptions in insulin receptor (IR) function occur in both disease states due to a decrease in insulin production in Type I Diabetes, and a decrease in insulin sensitivity with type II Diabetes. This conservation of altered insulin receptor signaling suggests an important role for IR in diabetic pathologies, including disrupted bone homeostasis. Insulin is a potent regulator of osteoblast function, with insulin stimulating collagen synthesis [13], and production of alkaline phosphatase [14]. Importantly, exogenous insulin can also reverse deficits in bone formation during healing in diabetic mice [9]. Several in vivo studies have examined the effects of IR conditional deletion in osteoblast-lineage cells [15-17], and demonstrate that IR deletion in osteoblasts impairs bone growth and homeostasis during skeletal maturation. However, the long-term effects of IR loss of function have not been studied, although there is high clinical relevance to this approach as increasing fracture risk is associated with increasing duration of Diabetes [18].

While the effects of IR deletion during different stages of osteoblast maturation are well understood, the effects of IR deletion in other bone-associated cells are unknown. S100a4 belongs to the S100 family of EF-hand Ca^2+^-binding proteins, which are expressed in highly cell- and tissue-specific patterns [19]. S100a4 is a potent negative regulator of osteoblast differentiation in vitro [20], with these effects mediated in part through inhibition of mineralization [21]. S100a4^-/-^ mice have larger bones, with increased bone density, and decreased osteoclastogenesis, while ShRNA knock-down of S100a4 increases cortical thickness [22]. S100a4 is expressed in perichondrium cells during bone development [23], and Duarte *et al*., demonstrated that S100a4 is expressed by pre-osteoblasts, followed by a decline in expression in terminally differentiated osteocytes, in vitro [20]. However, the expression pattern of S100a4 in adult bone is not clear. Here, we demonstrate that S100a4-lineage cells are found in cortical and trabecular bone, as well as the bone marrow. We generated S100a4-lineage specific IR conditional knockout mice and tested the hypothesis that conditional deletion of IRβ in *S100a4*-lineage cells would disrupt bone homeostasis leading to decreased mechanical properties. Loss of IR signaling in S100a4-lineage cells resulted in accelerated bone loss during aging and a significant reduction in torsional rigidity at 48-weeks of age, relative to WT littermates.

## Materials and Methods

### Ethics statement

All animal studies were approved and conducted in accordance with the University of Rochester University Committee on Animal Resources (UCAR) and the National Institutes of Health guide for the care and use of Laboratory animals.

### Animals

S100a4-Cre (#12641), IR^flox/flox^ (#6955)[24], Rosa-Ai9 (#7909)[25], S100a4^GFPPromoter^ (#12893)[26] mice were purchased from Jackson Laboratories (Bar Harbor, ME).

### Spatial localization of S100a4^+^ cells

To assess the spatial localization of S100a4-lineage cells and cells actively expressing S100a4, S100a4-Cre; Rosa-Ai9^F/+^ (S100a4^Ai9^) and S100a4^GFPPromoter^ mice were used, respectively. Male mice were sacrificed at 15 and 48 weeks of age and femurs were harvested and processed for cryosectioning. Briefly, femurs were fixed in 10% neutral buffered formalin (NBF) for 24hrs, decalcified for 3 days in 14% EDTA [27], immersed in 30% sucrose in PBS overnight and embedded in Cryomatrix. Serial 8μm sections were cut through the sagittal-plane of the femur and placed on Cryotape (Section-Lab, Japan). Sections were then stained with Hoescht 33342 (#R37605, ThermoFisher), coverslipped (Prolong Gold #P36930, Thermo Fisher, Waltham, MA), and digitally imaged using an Olympus slide scanner for endogenous Ai9 (TdTomato) and GFP fluorescence.

### Protein isolation and western blotting

Following sacrifice, femurs were harvested, stripped of surrounding soft tissue and the marrow was flushed. Total protein was harvested from the metaphysis via homogenization in RIPA buffer containing protease/ phosphatase inhibitors (#78445, ThermoFisher). Western Blots were probed with the following antibodies: Insulin Receptor β (1:100, #SC-20739, Santa Cruz), βactin (1:2500, #A2228, Sigma), Goat anti-rabbit secondary (1:2000, #1706515, BioRad). Blots were then imaged using West Pico (#34578) or Femto substrate (#34095, ThermoFisher).

### MicroCT analysis

Following sacrifice, the femur was isolated and cleaned of excess soft tissue. Femurs were imaged using VivaCT 40 cone-beam CT (Scanco Medical) and reconstructed in 3D at a resolution of 10μM (n=5-8 per genotype per age). Cortical bone was analyzed from the distal end of the third trochanter, where the lateral projection changes from a sharp point to a rounded point. This initial slice and the next 30 distal slices were included in the region of interest. Contours were drawn at the bone surface to exclude any soft tissue outside the bone. The threshold was set at 340, equivalent to a linear attenuation coefficient of 2.720 cm^-1^. Analysis of femur trabecular bone occurred from the proximal end of the growth plate proximally for 100 slices (10.5 micron slices). Contours were close-drawn to the cortical shell, and then shrunk to 95% in X and Y dimensions to avoid any inclusion of cortical bone. A threshold of 280 was used, which is equivalent to a linear attenuation coefficient of 2.240 cm^-1^.

### Biomechanical torsion testing

Following microCT, the ends of the femurs were cemented (Bosworth Company) in aluminum tube holders and tested using an EnduraTec TestBench™ system (Bose Corporation, Eden Prairie, MN). The femurs were tested in torsion until failure at a rate of 1deg/sec.

### Statistical analyses

Data are presented ± Standard Error of the Mean (SEM). For quantitative metrics, statistically significant differences were determined using two-way ANOVA with Tukey post-test. Differences were considered statistically significant at p<0.05.

## Results

### S100a4-lineage cells are found in trabecular bone osteoblasts and cortical bone osteocytes

To trace S100a4-lineage cells in bone, femurs were harvested at 15 and 48 weeks of age from male S100a4-Cre; Rosa-Ai9 mice, resulting in Ai9 (TdTomato) fluorescence in cells that have undergone Cre-mediated recombination driven by the S100a4 promoter. By 15 weeks of age, S100a4-lineage cells had become cortical osteocytes and trabecular lining osteoblasts, and S100a4-lineage cells were also observed in the bone marrow (Fig 1). Consistent with this, S100a4-lineage cells remained in the cortical bone, as well as trabecular lining cells, although there was a marked reduction in both trabecular bone and S100a4-lineage+ lining osteoblasts at 48 weeks of age (Fig. 1). To determine which cells actively express S100a4 at a given time, S100a4GFP^Promoter^ mice were used. At 15 weeks, a few S100a4GFP^Promoter^ cells were observed in the bone marrow, and periosteum, however, no S100a4GFP^Promoter^ cells were observed in cortical bone. In addition, S100a4GFP^Promoter^ cells were observed in trabecular bone (Fig. 2). Interestingly, at 48 weeks a few cortical osteocytes actively expressed S100a4, in contrast to 15 weeks. In addition, some S100a4GFP^Promoter^ cells were located in trabecular bone (Fig. 2).

**Figure 1.**
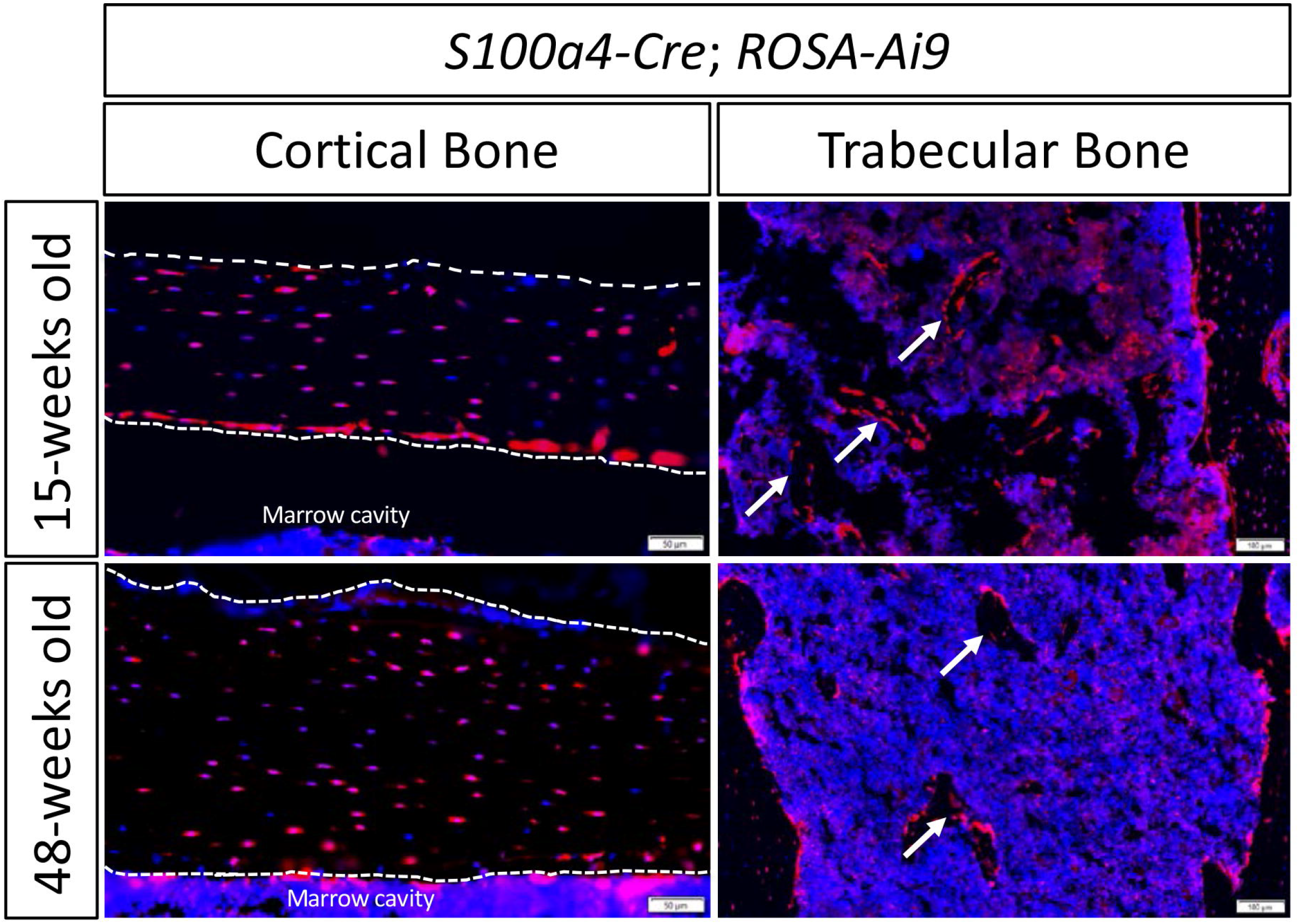
S100a4-lineage cells are located in cortical and trabecular bone. To determine the spatial localization of S1004-lineage cells in 15- and 48-week old femurs S100a4-Cre; Rosa-Ai9 mice were used. At 15- and 48-weeks of age S100a4-lineage cells were located in the cortical and trabecular bone, and the bone marrow. Scale bars represent either 50 or 100 microns as noted on each image.

**Figure 2.**
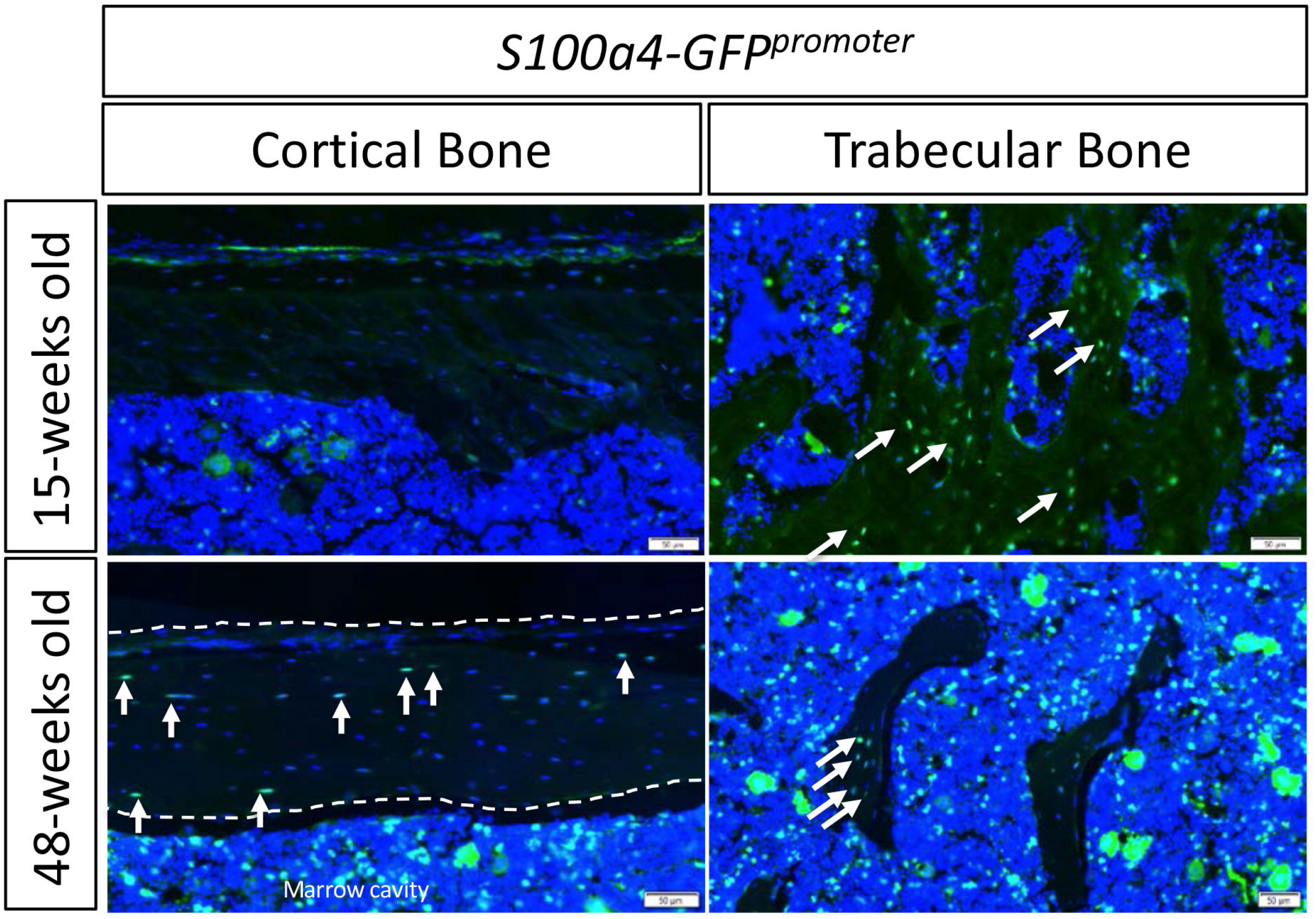
Cells actively expressing S100a4 are located in the trabecular bone, and cortical bone of aged 48-week but not 15-week old mice. To determine the spatial localization of cells actively expressing S100a4 at 15 and 48-weeks of age S100a4GFP^promoter^ were used. At 15 weeks of age S100a4+ cells were located in trabecular bone and the bone marrow, but not in the cortical bone. At 48-weeks S100a4^+^ cells were observed in cortical and trabecular bone, as well as the bone marrow. Scale bars represent 50 microns.

### IRcKO^S100a4^ decreases IR expression in bone

No changes in body weight, or fasting blood glucose were observed between groups [28] at 48 weeks of age, indicating that IRcKO^S100a4^ allows the delineation of IR function in S100a4-lineage cells independent of systemic metabolic changes. To confirm that IRcKO^S100a4^ effectively reduces IR expression in the bone, total protein was isolated from the metaphysis of WT and IRcKO^S100a4^ bones from 48-week-old animals. A substantial reduction in IRβ protein was observed in IRcKO^S100a4^ bones, relative to WT (Fig. 3), indicating efficient knock-down of IRβ in bone using S100a4-Cre.

**Figure 3.**
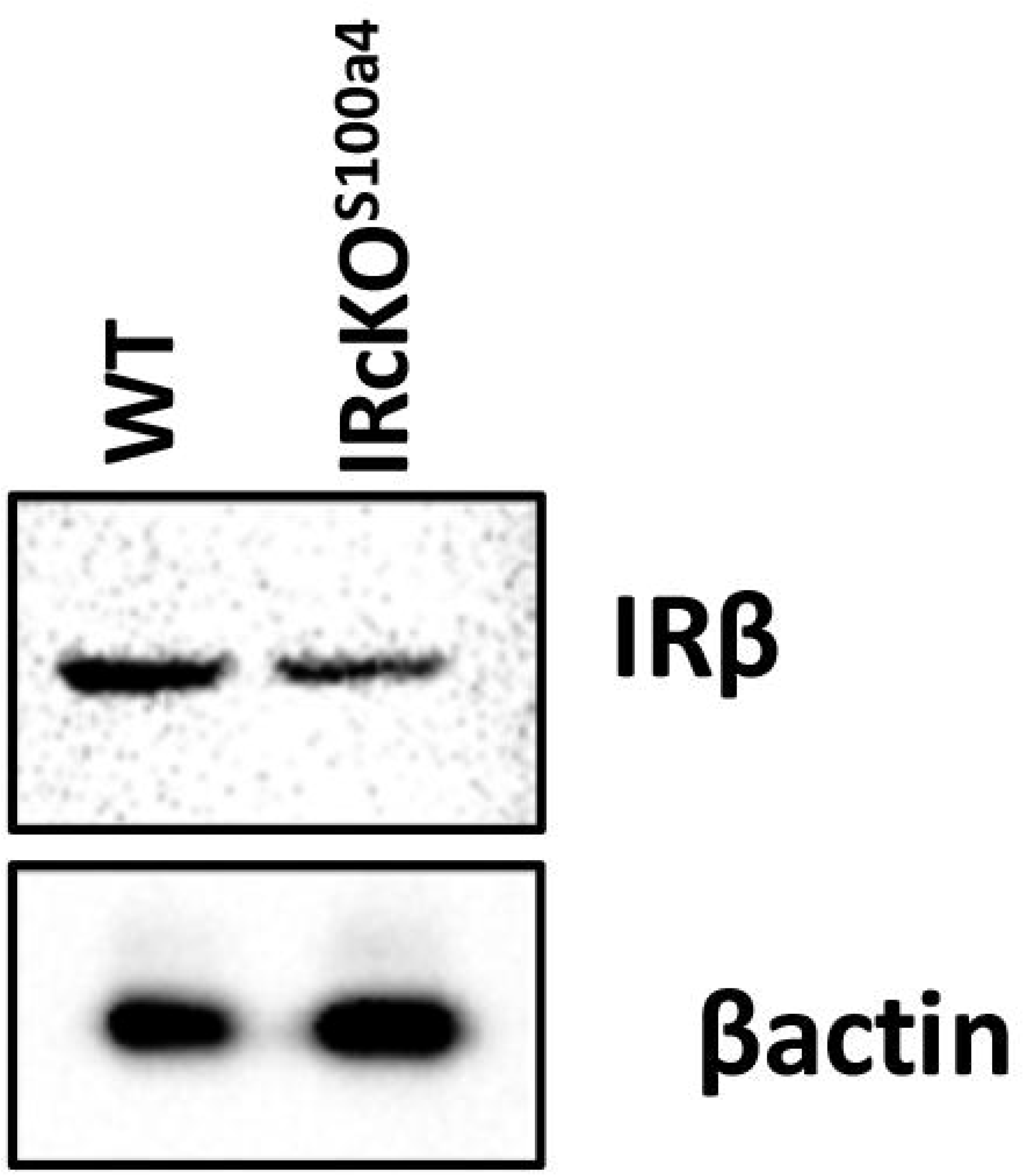
Deletion of IR in S100a4-lineage cells decreases IR expression in bone. At 48 weeks of age total protein was isolated from the femurs of WT and IRcKO^S100a4^ mice. Western blotting for IRβ demonstrates a substantial decrease in IRβ protein expression in IRcKO^S100a4^ relative to WT. Expression was normalized to βactin.

### IRcKO^S100a4^ enhances age-related trabecular bone loss

At 15 and 48 weeks of age no significant changes in Cortical Bone Volume were observed between WT and IRcKO^S100a4^ femurs (Fig. 4A-C). Trabecular BV was not different between WT and IRcKO^S100a4^ at 15 weeks of age (p=0.73). At 48 weeks Tb. BV was decreased in both WT and IRcKO^S100a4^ relative to their respective genotypes at 15 weeks, indicative of age-related bone less. In addition, Tb. BV. was significantly decreased in IRcKO^S100a4^ femurs relative to WT at 48 weeks (WT: 0.188mm3 ± 0.017; IRcKO^S100a4^: 0.048 ± 0.009, p=0.002) (Fig. 4D). Consistent with this, Tb. BV/TV was decreased in both genotypes at 48 weeks, relative to their respective genotypes at 15-weeks (P<0.0001). In addition, Tb. BV/TV was significantly decreased in IRcKO^S100a4^ at 48 weeks, relative to age-matched WT femurs (WT: 0.073 ± 0.007; IRcKO^S100a4^: 0.026 ±0.0023, p=0.0024) (Fig. 4E). While Tb. N (WT: 9% decrease, p<0.0001; IRcKO^S100a4^: 12% decrease, p<0.0001), and Tb.Sp (WT: +99%, p=0.0018, IRcKO^S100a4^: +141%, p<0.0001) were significantly altered as a function of age in both WT and IRcKO^S100a4^ animals, no differences were observed between genotypes at either time-point for any of these parameters (Fig. 4F & G). No differences in Trabecular Thickness were observed as a function of age or genotype (Fig. 4H)

**Figure 4.**
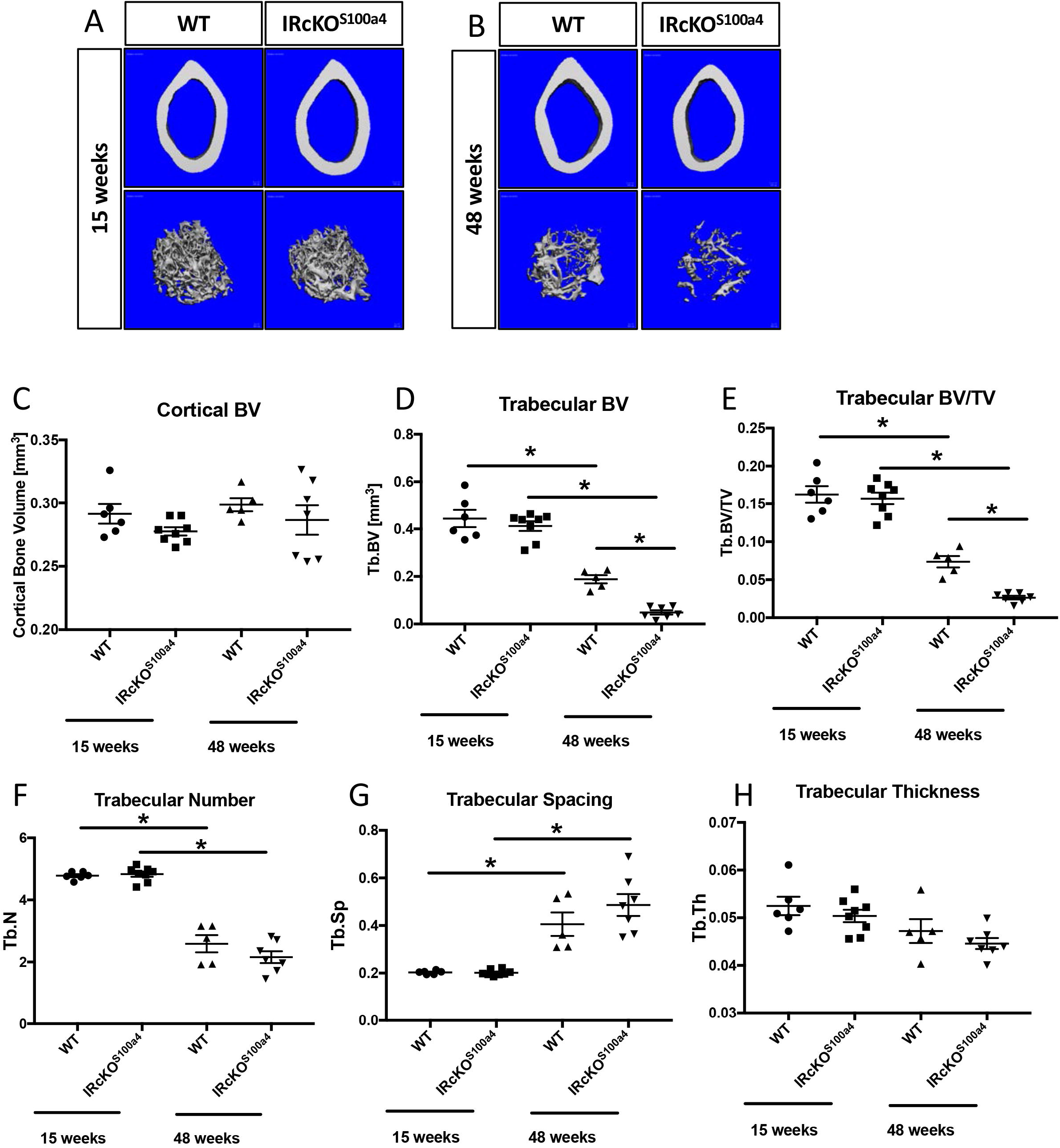
IRcKO^S100a4^ enhances trabecular bone volume loss during aging. (A & B) 3D microCT reconstructions of the cortical and trabecular bone of the femurs from (A) 15-week old, and (B) 48 week old male mice. (C-H) MicroCT analysis of (C) Cortical Bone Volume, (D) Trabecular Bone Volume, (E) Trabecular Bone Volume/ Total Volume, (F) Trabecular Number, (G) Trabecular Thickness, (H) Trabecular Spacing from WT and IRcKO^S100a4^ at 15 and 48 weeks of age. (*) indicates p<0.05.

### IRcKO^S100a4^ decreases torsional rigidity in aged mice

No changes in ultimate torque were observed between WT and IRcKO^S100a4^ femurs at 15 or 48-weeks age (Fig. 5A). In addition, no differences in Torsional Rigidity (T.R.) were observed between WT and IRcKO^S100a4^ femurs at 15 weeks of age (Fig. 5B), and T.R. was not significantly different between 15 and 48-week old WT mice (p=0.91). However, a significant reduction in T.R. was observed in 48-week-old IRcKO^S100a4^ femurs, relative to both 48-week old WT (26% decrease, p=0.02) and 15-week old IRcKO^S100a4^ (33% decrease, p=0.008).

**Figure 5.**
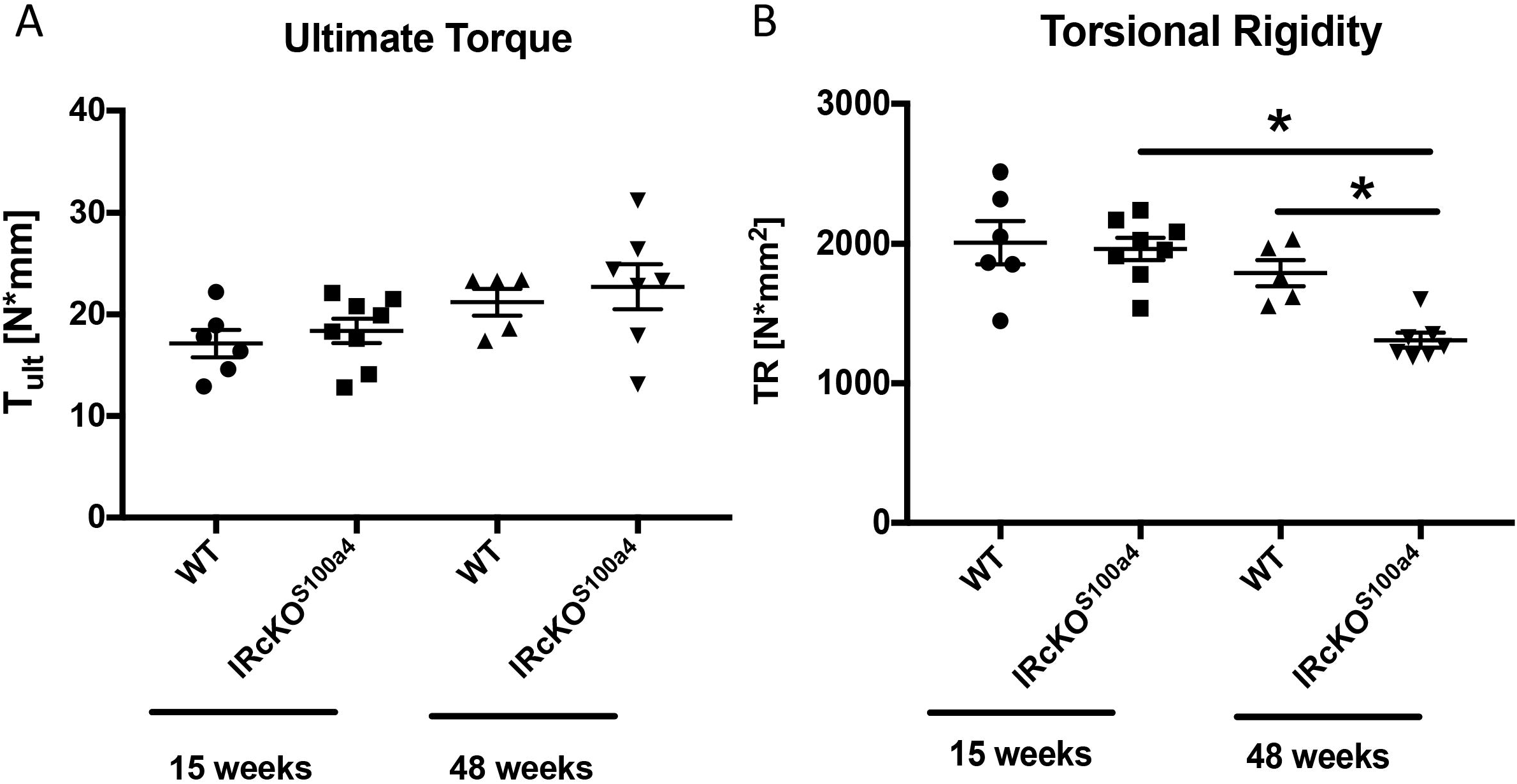
IRcKO^S100a4^ decreases femur torsional rigidity at 48-weeks of age. To determine the effects of IRcKO^S100a4^ on the torsional mechanical properties, femurs from WT and IRcKO^S100a4^ littermates underwent torsion testing at 15- and 48-weeks of age to assess A) Ultimate Torque at Failure, and B) Torsional Rigidity. (*) indicates p<0.05.

## Discussion

Type I and Type II Diabetes dramatically impair bone health, leading to a substantial increase in fracture risk [8, 29]. Previous work has demonstrated that bone is an insulin target tissue, with osteoblasts expressing functional insulin receptors, and insulin exerting bone anabolic effects[13]. Several studies have assessed the effects of IR deletion in distinct bone cell populations and demonstrate disruptions in bone homeostasis [15-17]. In the present study we have used the S100a4-Cre driver and demonstrate effective targeting of bone, and efficient deletion of IRβ in IRcKO^S100a4^. Loss of IRβ in S100a4-lineage cells results in bones that are more susceptible to age-related declines in trabecular bone, and decreased torsional rigidity, relative to age-matched WT mice.

While models of Type I and Type II diabetes clearly demonstrate the impact of systemic alterations in IR signaling on bone, a series of elegant studies have defined the effects of IR deletion specifically in bone cells. Fulzele *et al*., demonstrated the IR was expressed in trabecular bone osteoblasts, and that IR deletion using Osteocalcin-Cre (Oc-Cre) resulted in impaired bone acquisition. Deficits in trabecular bone were especially pronounced at 3 and 6-weeks of age, while minimal differences were observed between genotypes at 12-weeks of age [16]. Deletion of IR using Col1a1-Cre also disrupted initial bone formation with decreases in bone volume fraction and bone formation rate observed at 8 weeks of age, relative to WT [15]. Finally, Thrailkill *et al*., utilized Osterix-Cre (Osx-Cre) to drive IR deletion in osteoprogenitors, which resulted in decreased trabecular bone volume and thickness at 12-weeks of age [17]. Interestingly, we did not observe any alterations in initial bone acquisition (15-weeks of age) in IRcKO^S100a4^ mice, while dramatic impairments in bone maintenance were observed at 48-weeks of age. However, bone phenotypes were not assessed past 12-weeks of age in previous studies, so it is unknown how loss of IR in these populations may alter bone homeostasis during aging. Importantly, deletion of IR using both Col1a1-Cre and OC-Cre resulted in altered systemic metabolism, including increased adiposity and body weight [16], and altered glucose tolerance [15, 16]. In contrast, IR deletion in S100a4-lineage cells does not alter metabolism or induce obesity, even up to 48-weeks of age. Thus, IRcKO^S100a4^ permits assessment of IR function in this cell population without the confounding effects of altered systemic metabolism.

The difference in temporal bone phenotypes between IR deletion in osteoblast-lineage cells and S100a4-lineage cells may suggest a potential shift in the cell population(s) that rely on IR to maintain bone homeostasis over the course of aging, with a greater reliance of S100a4-lineage with increasing age. However, the identity of S100a4 cells with respect to the osteoblast lineage remains unclear. In the present study we have conducted traces of both S100a4-lineage cells and cells actively expressing S100a4 in adult mice. In contrast to previous studies [30], we show that both S100a4-lineage cells and a few actively expressing S100a4 cells are observed in cortical bone osteocytes, as well as in trabecular bone osteoblasts. Consistent with this, Duarte *et al*., observed that S100a4 was expressed by pre-osteoblasts, followed by a decline in expression in terminally differentiated osteocytes, in vitro [20], while Inubushi *et al*., demonstrated efficient targeting of the perichondrium during bone development using the S100a4-Cre [23]. These data suggest that IRcKO^S100a4^ results in deletion of IRβ in osteoblast precursors and a subset of mature osteoblasts, although the differences in phenotype compared to IR deletion in osteoblast lineages suggest that S100a4-cells may represent a potential discreet subpopulation of these cells.

There are several limitations to this study. First, the S100a4-Cre is non-inducible. Thus, we do not know the effects of delayed knock-down of IRβ in S100a4 cells. However, we have examined the bone phenotype in both young and aged mice, and do not observe any differences between genotypes in young mice. In addition, while S100a4-lineage cells are observed in trabecular bone and osteoblasts/ osteocytes, and S100a4GFP^Promoter^ expression confirms active expression of S100a4 in these cells, it is not yet clear how S100a4 relates to other bone cell populations. Thus, future work will be needed to more clearly understand the lineage, fate and function of S100a4 cells in bone. Finally, we have only investigated the bone phenotypes in male mice, as these mice were obtained as part of separate study. Given the dramatic reduction in bone in female mice with aging, it will be important to understand how disruption of IRβ expression and signaling in S100a4-lineage cells may alter this phenotype in female mice. Finally, we only assessed the effects of IRcKO^S100a4^ on the femur, although other locations such as the Lumbar spine are also sensitive and susceptible to age-induced bone loss.

Taken together, these data demonstrate that deletion of IRβ in S100a4-lineage cells is sufficient to accelerate age-related trabecular bone loss and impair mechanics. This study defines the importance of determining how IR signaling in S100a4-lineage cells alters osteoblast, osteoclast and osteocyte function with the goal developing therapeutic approaches to prevent impaired IR signaling induced bone loss.

## Acknowledgements

We would like to thank the Histology, Biochemistry and Molecular Imaging (HBMI) and the Biomechanics, Biomaterials and Multimodal Tissue Imaging (BBMTI) Cores for technical assistance. Research reported in this publication was supported by the National Institute of Arthritis and Musculoskeletal and Skin Diseases of the National Institutes of Health under Award Numbers K01AR068386, R01AR073169 and P30AR069655. The content is solely the responsibility of the authors and does not necessarily represent the official views of the National Institutes of Health.

**Author Contributions:** Study conception and design: AEL; Acquisition of data: VS, EK; Analysis and interpretation of data: VS, EK, AEL; Drafting of manuscript: AEL; Revision and approval of manuscript: VS, EK, AEL.

